# Fasta-O-Matic: a tool to sanity check and if needed reformat FASTA files

**DOI:** 10.1101/024448

**Authors:** Jennifer M Shelton, Susan J Brown

## Abstract

**Background:** As the sheer volume of bioinformatic sequence data increases, the only way to take advantage of this content is to more completely automate robust analysis workflows. Analysis bottlenecks are often mundane and overlooked processing steps. Idiosyncrasies in reading and/or writing bioinformatics file formats can halt or impair analysis workflows by interfering with the transfer of data from one informatics tools to another.

**Results:** Fasta-O-Matic automates handling of common but minor format issues that otherwise may halt pipelines. The need for automation must be balanced by the need for manual confirmation that any formatting error is actually minor rather than indicative of a corrupt data file. To that end Fasta-O-Matic reports any issues detected to the user with optionally color coded and quiet or verbose logs.

Fasta-O-Matic can be used as a general pre-processing tool in bioinformatics workflows (e.g. to automatically wrap FASTA files so that they can be read by BioPerl). It was also developed as a sanity check for bioinformatic core facilities that tend to repeat common analysis steps on FASTA files received from disparate sources. Fasta-O-Matic can be set with format requirements specific to downstream tools as a first step in a larger analysis workflow.

**Availability:** Fasta-O-Matic is available free of charge to academic and non-profit institutions on GitHub.

## Background

Sequence data can be stored as text with each letter representing a nucleic acid (DNA and RNA) or amino acid (protein). The linear nature of these molecules makes it natural to represent them as strings, finite sequences of characters. Although it has been argued that a graph, a network of edges connected by vertices, is a more accurate way to store genomic sequences because graphs allow the inclusion of alternate alleles and alternate possible assemblies [1] all of the most common methods for storing sequences (FASTA, FASTQ, SAM/BAM) use a linear strings.

Other decisions about how to represent sequence data can be more arbitrary. For example, any character that is not used as a base or an amino acid could be used to indicate the beginning of a new sequence. Additionally text could be wrapped to limit the information content in any one line of a file. The advantage of wrapping text is that some programs can then be designed to work one line at time limiting the burden of each step (e.g. the program would never have to process an entire chromosome of sequence data in a single step). The disadvantage is that code must be slightly more complex to load an entire sequence record into the working memory.

### 0.1 FASTA file format specifications versus recommendations

The popular FASTA file format stores sequence records and has very minimal format requirements [2]. Each sequence is preceded by a header/description line that begins with a > symbol. Sequence lines can include any standard International Union of Pure and Applied Chemistry (IUPAC) single character symbols for nucleic acids or amino acids or the ambiguous codes that indicate possible residues or bases [3]. They can also include - to indicate alignment gaps and * to indicate stop codons.

NCBI recommends wrapping FASTA file sequences lines [2]. It is also common practice to use the first ‘word’ in a header (i.e. any character string to the left of the first space in the header) as the unique sequence id. Although these features are common they are not required leading to format compatibility issues with tools that treat these conventions as required.

### 0.2 Customizing FASTA files to ensure that information is properly interpreted by downstream tools

Regardless of whether a FASTA file is technically improperly formatted or it’s format merely violates a popular convention, it is critical to quality analysis workflows that data is converted into a format that will be correctly interpreted by down-stream tools. Formatting issues can fall into multiple categories including actual format errors and formats that are not technically wrong but are non-standard, causing some tools to throw an error.

Some format errors indicate a major problem like an attempt to use the wrong data format (e.g. the first line is not a FASTA header because it does not begin with a > character). These types of errors will be subsequently referred to as fatal. Alternately, some formatting issues occur commonly without indicating the FASTA file is corrupt (e.g. improperly wrapped/unwrapped sequence lines, missing final new line characters, unusual new line characters like \r). These issues will be referred to as non-fatal. Fatal formatting issues should cause processing to stop. Non-fatal formatting issues should be automatically corrected according to the most common resolution for this type of error. While downstream processing continues, the analyst can double check the automated decision to reformat non-fatal issues. This way workflow would not be slowed for trivial reformatting steps and the more rare problems (e.g. when a missing last new line was caused by incomplete file transfer) could still be caught.

### 0.3 Existing tools

Existing bioinformatics tools address FASTA format inconsistencies. However many tools either halt and exit with an error (e.g. BioPerl [4], [5], [6]) or can produce reformatted output FASTA but cannot determine if there is a formatting issue to begin with (e.g. EMBOSS Seqret [7]).

The BioPerl module DB::Fasta will halt if a FASTA is inconsistently wrapped or if a line of sequence is too long (as in an unwrapped genome FASTA). This has the disadvantage of requiring human intervention to wrap and restart analysis.

Code:

~~~
#!/usr/bin/perl
~~~

~~~
use Bio::Seq;
~~~

~~~
use Bio::SeqIO;
~~~

~~~
use Bio::DB::Fasta; #makes a searchable db from FASTA file
~~~

~~~
my $out_file_temp = ’/home/bionano/test_db/all.fa’;
~~~

~~~
#Create new FASTA outfile object
~~~

~~~
my $seq_out = Bio::SeqIO->new(’-file’ => “>$out_file_temp”,’-format’ => ’fasta’);
~~~

~~~
#Load FASTA file as DB
~~~

~~~
my $db = Bio::DB::Fasta->new(“/home/bionano/test_db/miswrapped.fa”);
~~~

~~~
my $seq_obj = $db->get_Seq_by_id(’seq’); # get FASTA records using headers
~~~

~~~
#(where header = first ’word’ so really header whitespace should also be
~~~

~~~
#removed for this file)
~~~

~~~
$seq_out->write_seq($seq_obj);
~~~

Input:

~~~
>seq 1
~~~

~~~
ACTGTGTGCAATCGCTGNNNNCTCTCATCGGATCTTGCAATCGCTNNNCTCTCATCGGATTGCAATCGCTNNNCTtcatcCGGAT
~~~

~~~
CGCTGNNNNCTGTGTGCAATCGCTGNNNNCTCCTGATCGCTGNNNNCTGTGTGCAATCGCTGNNNNCTCCTGCAATCGCTGNNNN
~~~

~~~
CTCCTGTTCGNATCGatcctctgtttatgcttatagctagctgatcgtagnnntcaacgt
~~~

~~~
CTAGAGCGCAGCTCTGGGGGATTACTACTCACTACATCATTAGATCAGATacgactcann
~~~

~~~
>seq 2
~~~

~~~
cttatagctagctgatAATCGCTGNNTCATCGGATCTTGCCTTGCAATCGtcatcCGtcC
~~~

~~~
CGCTGNNNNCTGTGTGCAnnnnnnnnnnncgtaaaacgcctcctccgactcgTCTCTAGG
~~~

~~~
CTAGAGCGCAGCTCTGGGGGATTACTACTCACTACATCATTAGATCAGATacgactcann
~~~

~~~
nnnctacgCTATCAGGTCTCGAG
~~~

~~~
>seq 3
~~~

~~~
ATCAGCGCTCTATATGGCTCTGATTATAGTTTGCATTCATATGCTGATCTTctcagnntc
~~~

~~~
cttgacgctcgctATCTGTAGATCTGTACTtcagacagctcTCAGCAGNNNCTCAGCAGC
~~~

~~~
CTACGACAGTcatgcagactagcagt
~~~

Output:

~~~
------------- EXCEPTION -------------
~~~

~~~
MSG: Each line of the fasta entry must be the same length except the last. Line above #5 ’CTAGAGCGCAGCTCTGGGGG..’ is 61 != 86 chars…
~~~

EMBOSS seqret was designed as a very flexible tool to convert from one properly formatted file to another properly but distinctly formatted file. It also was designed to accept poorly formatted data (e.g. a FASTA missing the final new line that is improperly wrapped) and export a reformatted file (e.g. wrapped after 60 bases with a final new line).

Code:

~~~
seqret -stdout -sequence test.fa -outseq test_reformat.fa
~~~

Input:

~~~
>my header
~~~

~~~
AAAAAAAAAAAATTTTTTCCCCGGCGCGCGCGCTATAGCGCTATANNNNNNNNNNNNNNN
~~~

~~~
ATATATATATAT
~~~

~~~
ATTATTATATATATATTCTCTCTGGGCTCGCGTCTCGCTATTTATATATATATATATATTGCGCTCTCGTCTCCT
~~~

Output:

~~~
>my header
~~~

~~~
AAAAAAAAAAAATTTTTTCCCCGGCGCGCGCGCTATAGCGCTATANNNNNNNNNNNNNNN
~~~

~~~
ATATATATATATATTATTATATATATATTCTCTCTGGGCTCGCGTCTCGCTATTTATATA
~~~

~~~
TATATATATATTGCGCTCTCGTCTCCT
~~~

However, seqret does not log the detected errors in the format. Another feature of Seqret is that an output file is created even if the output is identical to the input. Storing two identical files is an inefficient use of disk space. Seqtk [8] is another example of a tool that can automate FASTA reformatting but does not first check original format or report format issues.

Another case to note is when an improperly formatted FASTA file is actually distributed as a component of a bioinformatics tool. Trimmomatic adapter sequences [9], for example, are distributed versions of the proprietary Illumina sequencing adapters but the FASTA files are missing final new lines. This can cause issues downstream if a workflow includes common analysis techniques like FASTA file concatenation.

The process of restarting analysis manually after wrapping a FASTA file may only take minutes. The time consuming aspect of this interruption is the time it takes the analyst to become available and the number of jobs this step must be repeated for. Likewise, storage of one extra FASTA file is trivial unless the FASTA file in question stores a whole genome in which case the burden can add up for a bioinformatics core. Efficiency and automation are crucial as bioinformatic analysis projects become more numerous and time consuming. Many tools can either detect a format issue or repair a format issue. No existing tool was found that both validates FASTA format and reformats automatically only where required for a user defined list of non-fatal FASTA format issues.

## 1 Implementation

Fasta-O-Matic was designed to fit seamlessly into an analysis workflow. It detects which format issues are actually present in the FASTA file and then only produces a reformatted file if the current file violates the user defined format requirements.

### 1.1 Portability

Where possible Fasta-O-Matic was designed to be easy to distribute and use. Fasta-O-Matic is distributed on GitHub under the MIT license to allow for easy access to or customization of the code. The tool was also built and tested on both Python2.7 and Python3.3 to minimize incompatibility with existing linux environments. The script generates complete help menus when called from the command line with the `--help` command and from within python with `help(fasta_o_matic)`. Additionally, Fasta-O-Matic includes a sample FASTA file with missing new lines, inconsistent wrapping and spaces in headers along with a tutorial which describes how to reformat the sample. These features ensure that Fasta-O-Matic is easy to incorporate into existing workflows.

### 1.2 Automate where appropriate

The script was designed to efficiently execute the most likely solution given the presence or absence of format issues. Fasta-O-Matic returns a filename for the output FASTA file that conforms to the user defined format. If the original file already conforms, then Fasta-O-Matic returns the original filename rather than outputting a redundant FASTA file under a new name.

Fasta-O-Matic will exit and report an error if the FASTA file cannot be read, the default or defined output directory cannot be written to, the input FASTA file does not begin with a > or if any sequence line includes a non-IUPAC character. The last two errors are considered to be fatal FASTA format errors.

Inconsistent or unwrapped sequence lines, spaces in headers and missing or non-standard new lines are considered non-fatal errors. Testing for these issues is optional. If they are detected, the decision is made to reformat as requested, report the issue to the analyst and continue the workflow.

Testing the uniqueness of the header/description line can return a non-fatal warning and a reformatted file or a fatal error. Testing for uniqueness is optional. If the first word in each header/description line is unique then it follows that all description lines are unique. If the first words are not unique then it is possible that is because the header ids include whitespace `‘>seq 1’` or `‘> seq 1’`. In this case a resolution is to replace the whitespace with a character. Fasta-O-Matic replaces the whitespace with an underscore and retests for the uniqueness of the first words in the headers. If this version passes than the user is warned that whitespace effected header uniqueness and was removed from headers. If removing whitespace also fails to resolve the issue the lack of uniqueness is considered a fatal error. The fatal error is reported and the program halts.

The script also automatically adjusts to run the minimal number of steps sufficient to fix and report format issues. If it is included in the set of quality control (QC) steps then wrapping is the first format issue tested because while repairing FASTA wrapping both headers and new lines can be corrected. New lines are given priority after wrapping because while repairing new lines it is also trivial to repair headers. Next, uniqueness of the header lines is tested. Finally, headers are evaluated for whitespace. If an early test returns a format issue and launches a reformatting that automatically repairs any remaining format issues then Fasta-O-Matic still tests for any additional format errors in the original file.

All format issues are reported in the programs logs in case they indicate an unexpected issue with the data. Logs can be optionally color coded so that red indicates errors, yellow indicates warnings (e.g. a non-fatal issue was found and automatically reformatted) and green indicates status information. This method of logging is designed to draw the attention of the bioinformatics analyst to relevant warnings or errors even if they have grown accustomed to seeing Fasta-O-Matic output frequently.

### 1.3 Workflow integration

Sequence FASTA files are often passed as arguments to commandline tools. For example FASTA files can be passed as an argument to bowtie2-build to be indexed as an alignment reference [10] or passed to trimmomatic as adapters to detect sequencing artifacts. The output filename used by Fast-O-Matic varies to reflect the reformatting performed. For seamless integration into automated workflows Fasta-O-Matic returns the full path of the new properly formatted FASTA file or the original file (if it is already formatted properly). This can be captured as a variable and used as an argument in subsequent commands. The Bash commands below show and example of capturing the FASTA file name as a variable.

Code (backslashes are used to indicate a new line that is for display in the article rather than the new lines being included in the actual code):

~~~
filename=“$(python fasta_o_matic.py -f NC_010473_mock_scaffolds.fna \
~~~

~~~
-o ∼/out_fasta_o_matic -c)”
~~~

~~~
echo $filename
~~~

## 2 Results

### 2.1 Data

FASTA format tools were tested on the Vicugna pacos-2.0.1 whole genome shotgun sequence scaffolds because the 2.17 Gb *Vicugna pacos* genome is large (> 1 Gb) and has many scaffolds (276727) [11]. The large genome size and high number of individual sequences should approximate a typical large FASTA file. The FASTA file was downloaded from the National Center for Biotechnology Information (NCBI) FTP as NW 005882702.1 *Vicugna pacos* isolate Carlotta (AHFN-0088) Vicugna pacos-2.0.1 assembly scaffolds. An additional unwrapped sequence was added to the end of the file. This sequence was also missing a new line. Each FASTA record in the file also had spaces within the text of the headers.

The additional simulated FASTA record is available on [GitHub](https://github.com/kstatebioinfo/Fasta-O-Matic-a-tool-to-sanity-check-and-if-needed-reformat-FASTA-files/blob/master/simulated_unwrapped.fa).

### 2.2 Reformatting tests

No tool was found with all of Fasta-O-Matic’s functions. Therefore sequence line wrapping was compared between Fasta-O-Matic and two other common reformatting tools, seqtk and seqret. Fasta-O-Matic was run with the `--qc_steps` flag set to either `wrap new_line header_whitespace unique (all), wrap (W) new_line` (NL), unique (U) or `header_whitespace (HW)`. Seqtk was run with the arguments `seq -l 60`. Seqret was run using only the `-sequence` and `-outseq` arguments. Code used in tests or to produce figures can be found on [GitHub](https://github.com/kstatebioinfo/Fasta-O-Matic-a-tool-to-sanity-check-and-if-needed-reformat-FASTA-files/tree/master/figures). Run time and max memory was reported for each tool. Tests were run on a Xeon Phi server with 48× 12-core Intel Xeon CPUs, 256GB of RAM, Linux CentOS 7 and Python2.7.

### 2.3 Comparison between results

All tools could reformat the improperly wrapped FASTA file. Fasta-O-Matic had the lowest maximum memory requirements (Figure 1, Table 1). This may be useful if working on a large genome on a local machine or cluster headnode where memory usage is restricted. Fasta-O-Matic took several minutes rather than seconds (seqtk and seqret took < 13 s) (Figure 2, Table 1).

**Figure 1.**
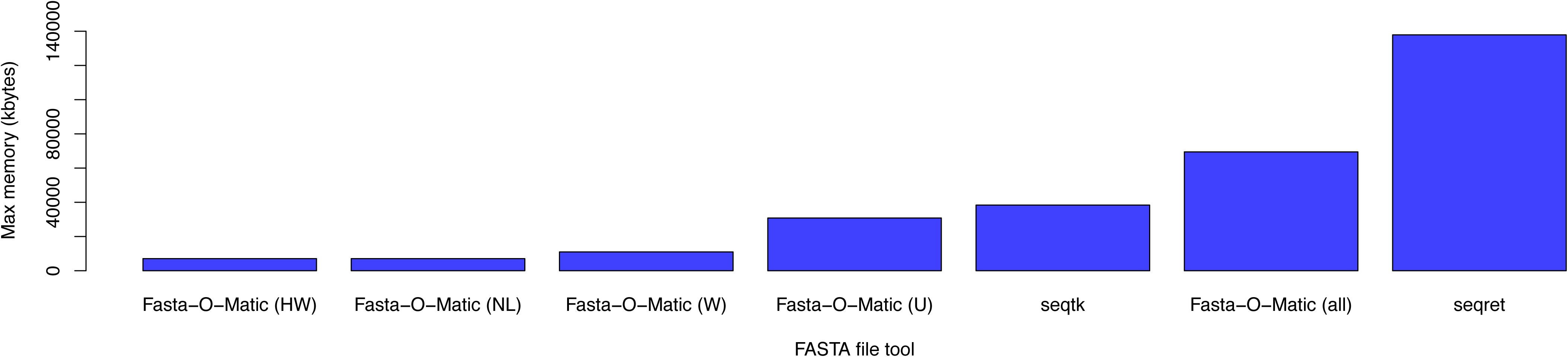
Max memory used by various FASTA tools. Tools were run on the *Vicugna pacos* isolate Carlotta (AHFN-0088) Vicugna pacos-2.0.1 whole genome shotgun sequence NW 005882702.1 with additional unwrapped FASTA sequence record.

**Table 1.**
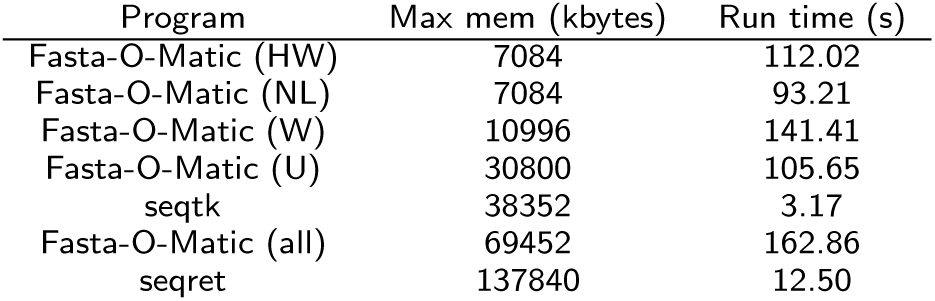
Runtime and max memory used by various FASTA tools. Tools were run on the Vicugna pacos isolate Carlotta (AHFN-0088) Vicugna pacos-2.0.1 whole genome shotgun sequence NW 005882702.1.

**Figure 2.**
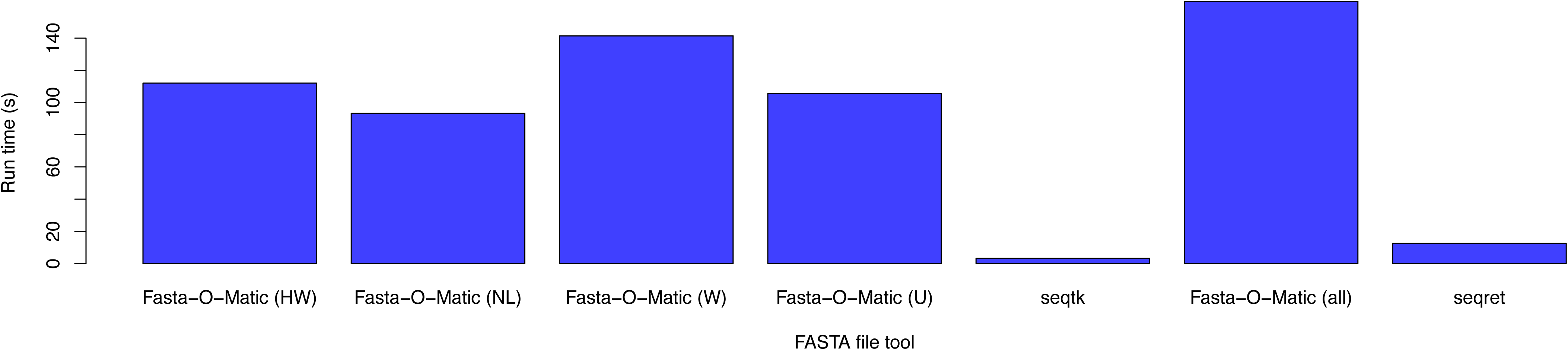
Run time for various FASTA tools. Tools were run on the *Vicugna pacos* isolate Carlotta (AHFN-0088) Vicugna pacos-2.0.1 whole genome shotgun sequence NW 005882702.1 with additional unwrapped FASTA sequence record.

Fully re-formatted simulated FASTA record (backslashes are used to indicate a new line that is for display in the article rather than the new lines being included in the actual FASTA record):

~~~
>NW_000000000.0 Vicugna pacos isolate Carlotta (AHFN-0088) FAKE genomic scaffold, \
~~~

~~~
Vicugna_pacos-2.0.1 Scaffold-, whole genome shotgun sequence
~~~

~~~
ATACAACCATAAAGGTGCTATTCAGTCCATGGTTACAGGACATAACTACAACACACACCC
~~~

~~~
ACGTACACATGCGCATGCGCATGCACACACCCACGTACACGTACACGTACGCATACACAC
~~~

~~~
CCACGTACACGTACACGTACGCATACACACCCACGTACACGTACACGTACGCATACACAC
~~~

~~~
CCACGTACACGTACACGTACGCATACACACCCACGTACACGTACACGTACGCATACACAC
~~~

~~~
CCACGTACGCACACACGTACACGTGTAGGCACGCATTTAGCAAGTATTTAGCTTGCTTAA
~~~

~~~
ACAAACCCCCCCTACCCCCCACGAGCCCCACCTTATATACCAGACAGTCTTGCCAAACCC
~~~

~~~
CAAAAACAAGACATAGCGCATAAGCTATAGAACCCGGACAAACCTTTGCCCACAAACCCA
~~~

~~~
ACTTCTTAAATAATCACATGGCCAAATCGTACCAATGTGTTACTCTAGTATATTAAAAAT
~~~

~~~
ATACAGACAGCTATCTCCCTAGATCCGCCAAAATTTTTAAAACAGAATTCAACAACCTTT
~~~

~~~
TTAATGGCACCCCCCCCCCCCATAAATGACC
~~~

## 3 Conclusions

Overall, both memory and run time requirements were small for all three programs. However, the extra minutes taken by Fasta-O-Matic to test for fatal and non-fatal format issues may prevent hours lost waiting for an analyst to manually restart analysis or worse discover that a file was corrupt only after analysis is complete. Fasta-O-Matic was also the only tool identified that skips reformatting if none is required balancing the need to prepare data to be properly interpreted by bioinformatics tools with the practical need to conserve disk space. Fasta-O-Matic is a portable and easy to use tool to facilitate bioinformatics analysis by automating FASTA file inspection in busy bioinformatics cores.

## 4 Availability and requirements

**Project name:** Fasta-O-Matic tool

**Project home page:** The Fasta-O-Matic script and tutorial are available at

https://github.com/i5K-KINBRE-script-share/read-cleaning-format-conversion/tree/master/KSU_bio

**Operating system(s):** Linux (tested on CentOS 7, Gentoo and Ubuntu).

**Programming language:** Python2.7+, Python3.3+

**License:** Tool and tutorial are available free of charge to academic and non-profit institutions.

**Any restrictions to use by non-academics:** Please contact authors for commercial use.

**Dependencies:** Fasta-O-Matic requires the python modules Colorer and general which are distributed in the same git repository.

## Abbreviations

IUPAC: International Union of Pure and Applied Chemistry
QC: quality control
NCBI: National Center for Biotechnology Information
W: wrap
NL: new_line
HW: header_whitespace
U: unique

## Author’s contributions

JMS wrote most of the code for Fasta-O-Matic. JMS and SJB did the writing. Both authors read and approved the final manuscript.

## Acknowledgements

Thanks to Sheldon McKay https://github.com/mckays630 for contributing to the editing of the Fast-O-Matic program.

This project was supported by an Institutional Development Award (IDeA) from the National Institute of General Medical Sciences of the National Institutes of Health under grant number P20 GM103418. The content is solely the responsibility of the authors and does not necessarily represent the official views of the National Institute of General Medical Sciences or the National Institutes of Health.

